# Football as foraging? Movements by individual players and whole teams exhibit Lévy walk dynamics

**DOI:** 10.1101/2024.06.11.598528

**Authors:** Ivan Shpurov, Tom Froese, Takashi Ikegami

## Abstract

Many organisms, ranging from modern humans to extinct species, exhibit movement patterns that can be described by Lévy walk dynamics. It has been demonstrated that such behavior enables optimal foraging when resource distribution is sparse. Here, we analyze a dataset of football player trajectories, recorded during the matches of the Japanese football league to elucidate the presence of statistical signatures of Lévy walks; such as the heavy-tailed distribution of distances traveled between significant turns and the characteristic superdiffusive behavior. We conjecture that the competitive environment of a football game leads to movement dynamics reminiscent of that observed in hunter-gathering populations and more broadly in any biological organisms foraging for resources, whose exact distribution is unknown to them. Apart from analyzing individual players’ movements, we investigate the dynamics of the whole team by studying the movements of its center of mass (team’s centroid). Remarkably, the trajectory of the centroid also exhibits Lévy walk properties, marking the first instance of such type of motion observed at the group level. Our work concludes with a comparative analysis of different teams and some discussion on the relevance of our findings to sports science and science more generally.

## Introduction

Play, broadly construed as an activity performed of free will for the individual’s pleasure, is widely considered deeply rooted in the human psyche. *Homo sapiens* are not the only mammals who engage in it and its influence on the formation of human culture and civilization can be debated but not neglected (***Huizinga, 2014***). The commercial and societal success of team sports and ball games brought about a stream of scientific works seeking to elucidate factors informative of the successful performance of individual players and teams as a whole. In our work, we perform a quantitative analysis of trajectories recorded during the matches of the Japanese football league. Although we use some of the methods from the standard toolbox of quantitative football analysis, our main focus is to perform the game analysis implementing insights from the field of animal movement analysis. Our conjecture is that individual player movements exhibit a fat-tailed step size distribution, which is indicative of Lévy walk dynamics.

Lévy walk is a type of random walk, with its key characteristic being that the distribution of step lengths ***S*** is drawn from a scale-free (power-law) distribution. Thus ***P***(***S***) ∝ ***S***^−*k*^ and *k* lies in rang 1 *< k <* 3. Particles exhibiting Lévy walk dynamics manifest superdiffusive behavior, spreading faster than Brownian walkers. The resultant trajectory possesses fractal properties, with clusters of short steps interspersed with much longer sprints.

It has been conjectured that Lévy dynamics possess several advantageous qualities to motile biological agents which seek to maximize their encounter rate with resources of some sort. Notably, it prevents the agent from revisiting already explored terrain, and combining numerous short steps with long leaps enables the balance of exploration and exploitation. A complete review of the relevant literature would be outside the scope of this paper; we refer the interested reader to books and reviews (***Reynolds, 2018; Viswanathan et al., 2011***). To briefly outline the landmarks works, the first quantitative evidence dates back to the (***Viswanathan et al., 1996***), when Lévy walk behavior was discovered in the movement patterns of wandering albatrosses. In a curious turn of events, the statistical analysis of this seminal paper was found to be inadequate(***Edwards et al., 2007***). However subsequent work, relying on improved statistical techniques showed that wandering albatrosses exhibit Lévy walk dynamics (***Humphries et al., 2013***). This story is not only a curious incident in the annals of science history, it underlines the difficulties of distinguishing between the Lévy walk and competing hypothesis, a topic which will be expounded further on in our work.

Heavy-tailed step size distribution has been found in the diving pattern of aquatic predators (***Sims et al., 2008***), termite trajectories(***Miramontes et al., 2014***), T-cell movement (***Harris et al., 2012***), airborne seed dispersal (***Reynolds, 2013***), moving patterns of different types of terrestrial animals (***Ríos-Uzeda et al., 2019; Ramos-Fernández et al., 2004***), human mobility patterns inferred from cellphone data (***Gonzalez et al., 2008***) and foraging patterns of individuals in hunter-gather populations (***Reynolds et al., 2018***). It is worth noting that there is evidence to suggest that such movement pattern is evolutionarily ancient, as it is evidenced by the discovery (***Sims et al., 2014***) of Lévy walks in the fossilized trails of sea urchins, which are around 50 million years old. Currently, much of the research, concerning Lévy movement patterns in animal foraging, accepts its *de facto* presence, while the main focus is on elucidating its generative mechanism, with hypothesizes including collective effects (***Reynolds and Ouellette, 2016***), interactions between the foraging agent and its environment (***de Jager et al., 2011; Sims et al., 2008***) and inherent neural generators (***Sims et al., 2019***).

Concurrently with the exploration of animal foraging strategies quantitative analysis of football player movements has also blossomed in recent years (***Sarmento et al., 2018***). Several main venue of analysis emerged - examining the behavior of teams’ center of mass (centroid) (***Frencken et al., 2012, 2011***), investigating correlations between player pairs (***Marcelino et al., 2020***), and applying graph theoretic measures to complex networks derived from players’ interaction (***Cotta et al., 2013; Buldú et al., 2019***). Summarily, these works demonstrate that football teams exhibit a high level of cohesion and the motion of individual players is highly correlated, with their mates as well as with their opponents and centroid position, as well as player dispersion are informative of the games’ dynamics. This research remains mostly confined to the sports science field, with virtually no intersection with animal movement ecology and other quantitative behavioral science endeavors. We hope to address this discrepancy in our current work.

## Results

We use high-resolution (25 fps, centimeter precision) trajectories of players recorded during the matches of the Japanese Football League (J-league) during the 2022 season. Data was acquired from the league’s official data company, DataStadium, Inc. The dataset contains all games played by that season’s champions Hokkaido Consadole Sapporo. A full list of the matches contained in the dataset is provided in the supplementary materials. In this work the study properties of player’s trajectory, most notably their step-size distribution, both for individual players and for the whole teams. The rest of this paper is structured as follows: in the first section, we analyze single-match data while in the second section, we present summary statistics computed using data for all games in the dataset.

### Analysis of a Single match

For the illustrative analysis presented below, we choose a game played between Hokkaido Consadole Sapporo and Nagoya Grampus, on the 30th of July 2022. The game was played on the pith of dimension 105 × 68, which is standard for J-league. The aforementioned teams are hereafter referred to as Team 1 and Team 2. This game ended in a draw with a score of 2:2. Trajectories of all players for both teams have been used for analysis, whenever the player was present for the whole duration of the game or was substituted during its course. Both teams substituted players in the second half of the game, with team 1 substituting 3 players and team 2, 4 players.

Smothering with a Gaussian kernel with *σ* = 3 is applied separately for the x and y components of the trajectories to remove possible recording artifacts. Sensitivity analysis was conducted to show that varying *σ* within reasonable bounds does not alter the results. To obtain a systems’ level description of the teams’ activity we computed the center of mass trajectory, for each team, as 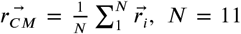. To ascertain to what degree players tend to cluster during the game we computed the mean distance of players from the center of mass of their team 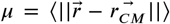 and its standard deviation *σ* as functions of time. These results are presented in Figure 1. Note, that both those quantities exhibit significant variability over time suggesting alternation of contraction and relaxation dictated by the dynamics of the game. Positions of the centers of masses for competing teams are visibly correlated.

**Figure 1.**
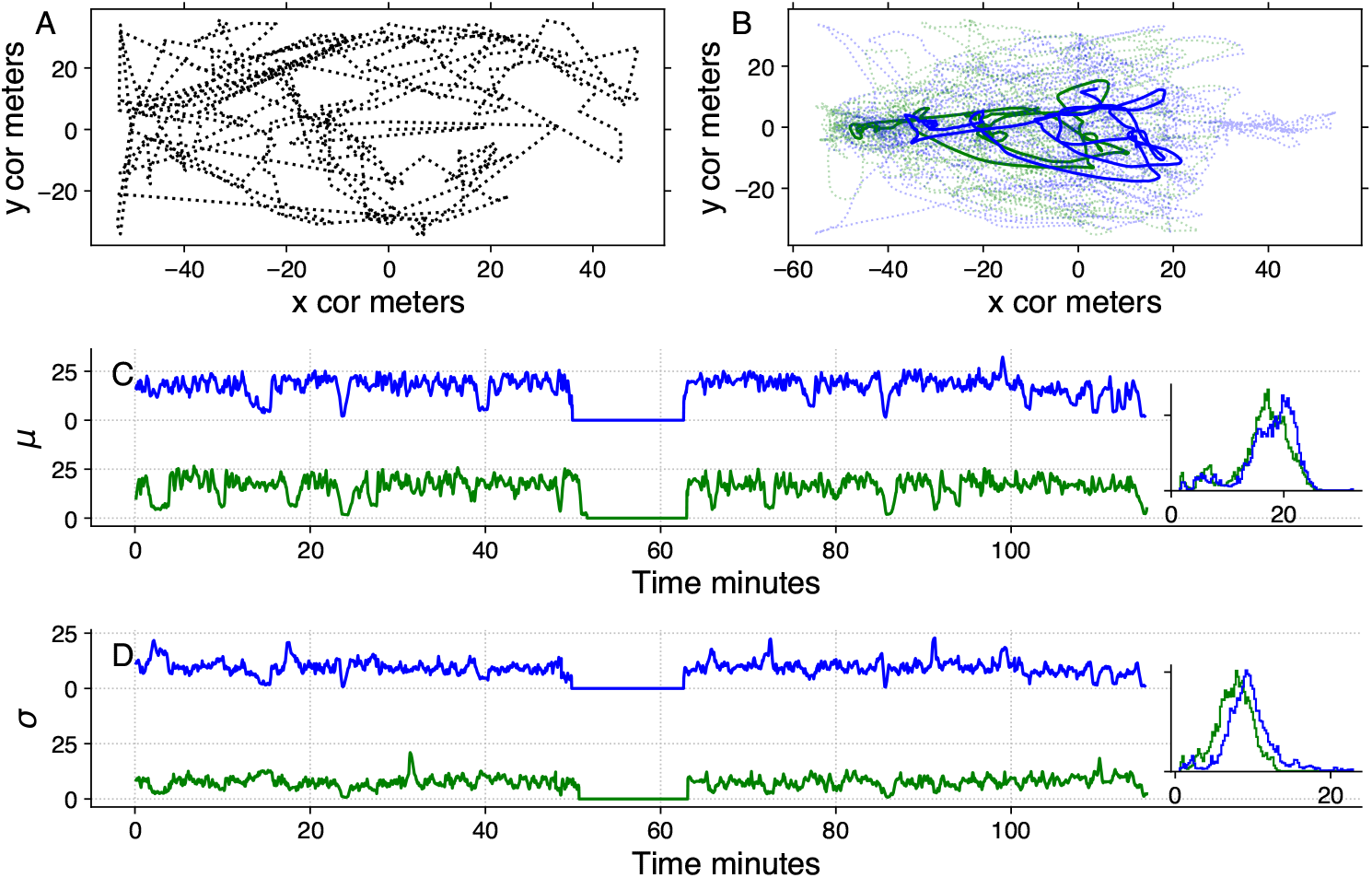
Pane A shows the ball’s trajectory during the first 5 minutes of the game. Pane B presents the trajectories of all players from both teams depicted with semitransparent dotted lines and trajectories of their respective centers of masses shown with solid lines. The first 5 minutes of the game are shown. The green color identifies Team 1, and the blue color identifies Team 2. Pane C depicts the mean distance *μ* of the player from their team’s center of mass for the game’s duration, while pane D shows the standard deviation from the center of mass position *σ*. Inset plots in the right corner of panes B and D show normalized histograms of *μ* and *σ* respectively.

First insights into the nature of biological movement can be often obtained from computing the Mean Square Displacement (MSD)(1). Due to the small (N=11 per team) number of agents in the field simultaneously, we used methods to approximate MSD from a single trajectory (***Michalet, 2010***). For each player, we partitioned the entire trajectory of its movements into 20-second intervals and computed the squared average displacement.

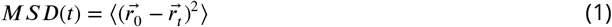

As it is known from statistical mechanics (***Berg, 1993; Viswanathan et al., 2011***) the exponent *a* relating the square of the displacement and time is informative of the type of motion: while in the case of Brownian motion, the variance of displacement is directly proportional to time, *a* = 1 higher or lower values indicate superdiffusive or subdiffusive motion respectively. Lévy walks are superdiffusive, therefore their exponents lie in the range 1 *< a <* 2, between the Brownian motion and the ballistic limit. As the third column of Figure 2 shows, values of *a* computed for individual players show that their motion is superdiffusive.

**Figure 2.**
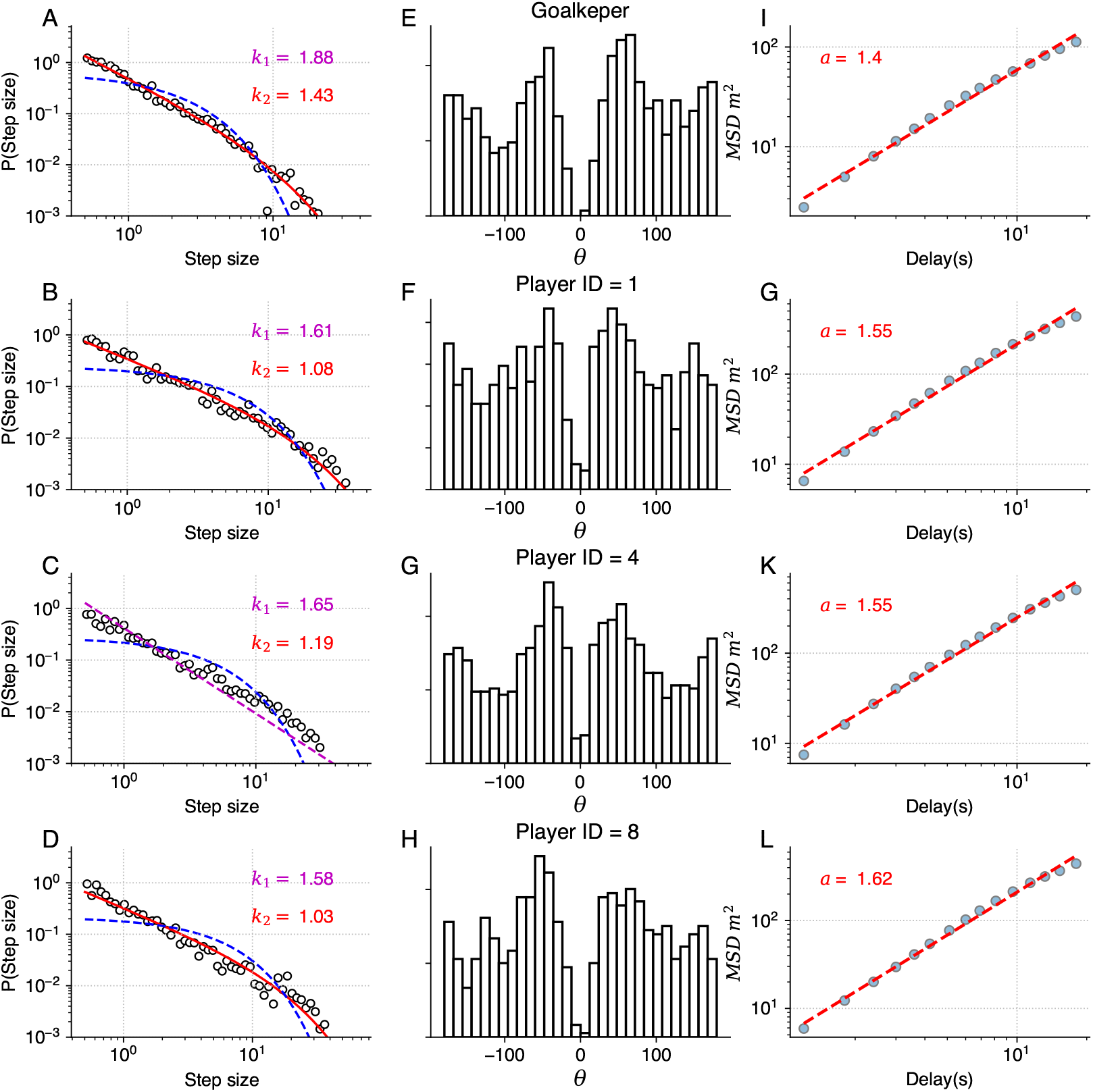
Analysis of four players from team 1. Panes A, B, C, and D demonstrate the step size distribution. The solid red line corresponds to the truncated power-law fit, the magenta line shows the power-law fit and the dashed blue line shows the exponential distribution. For visual clarity, power-law fit is only demonstrated on pane C, other panes show truncated power-laws. Exponents for both fits are given. Hollow black circles represent the normalized distribution histogram; logarithmic binning is used. Panes E, F, G, and H show the distribution of turning angles. Panes I, G, K, and L present the time *vs* MSD relation. The dashed red line shows the linear fit to the data.

Super diffusive behavior is a crucial characteristic of the Lévy walk, however, it can exhibited by other modes of motion. For the definitive test, it is necessary to examine the step-size distribution. As the players’ trajectories are continuous and two-dimensional, we implement the methodology developed by (***Humphries et al., 2013***) for such scenarios. This approach uses the symmetry of Lévy walks: when projected to a lower dimension Lévy walk retains its properties and remains distinguishable from other modes of movement, such as Brownian motion. Considering the 1D component of trajectory significant turns can be unequivocally determined as reversions of motion in one dimension. Motion in X and Y dimensions are considered separately, as we assume that the rectangular field would impose different constraints on player mobility in vertical and horizontal directions and therefore different values of Lévy walk exponents. Summary statistics for all players, including their Lévy exponents in both X and Y dimensions are available in the supplementary material. Different Lévy exponents for different axes of motion have been previously reported for airborne dispersal of seeds (***Reynolds, 2013***), concerning horizontal and vertical dimensions.

The first column of Figure 2 presents the distribution of step sizes for the Y dimension for the goalkeeper (Pane A) and 3 field players with different roles (remaining panes). We used the Maximum-likelihood estimator(MLE) to fit the power-law and truncated power-law to the data (***Alstott et al., 2014***) as well as to conduct log-likelihood tests to compare truncated power-law with other candidate long-tailed distributions, such as log-normal and exponential (Figure 2). For all players, these tests have shown unambiguously (p»0.05) that the truncated power-law is the best fit for the data. Power-law without truncation gives a better fit than the exponential distribution but is superseded by both the log-normal distribution and the truncated power law. Power-law scaling extends more than two orders of magnitude in the Y-axis and is close to two orders of magnitude on the X-axis, therefore being close to fulfilling the proposed (***Stumpf and Porter, 2012***) “rule on thumb” for elucidating scale-free phenomena in biological systems. The curtailed scale on the X-axis is caused by the size of the football pith. Values of *k*_1_ and *k*_2_ correspond to the Lévy walk exponent of regular and truncated power-laws; they are different for different players; most pronounced is the distinction between the goalkeeper and the field players. As is expected, power-law exponents *k*_1_ and *k*_2_ are inversely related to the MSD exponent *a*.

Another characteristic of the true Lévy walk is that the distribution of turning angles between consecutive steps is close to uniform. We use the points previously identified as reversals of 1D motion in the Y dimension to compute the angle direction of movement before and compute players’ turning angles at these points. The turning angle distribution is presented in the central column of Figure 2. Observed distributions could be interpreted as follows: small turning angles are rarely encountered, as the nature of the projection operation eliminated them from data, for *θ >* 0 the distribution is close to uniform. Uniform turning angle distribution distinguishes Lévy walk from the correlated random walk, which also has superdiffusive properties. Such turning angle distribution has been previously reported for marsupials (***Ríos-Uzeda et al., 2019***) and wandering albatrosses (***Humphries et al., 2013***). Both these species have been found to exhibit Lévy walk behavior with their step size distribution best approximated by truncated power-law.

Additionally, we study the distribution of the step sizes for movements of the centers of masses of both teams. We compute the center of mass positions as described previously and then compute the step size distributions for its movements in Y-dimension. We construct surrogate data as a control by performing a circular random shift of individual players’ position timieseries before computing the team’s center of mass position. These results are presented in Figure 3. An inspection of data presented on logarithmic and linear axes (inset) yields that longer steps are completely absent from the surrogate data. Max-likelihood fits prove that the step size distribution of the real data is best fitted with the truncated power-law distribution (*p <<* 0.05) while for synthetic data exponential distribution presents itself as the more likely candidate, although no clear distinction can be made using log-likelihood tests (*p >* 0.05).

**Figure 3.**
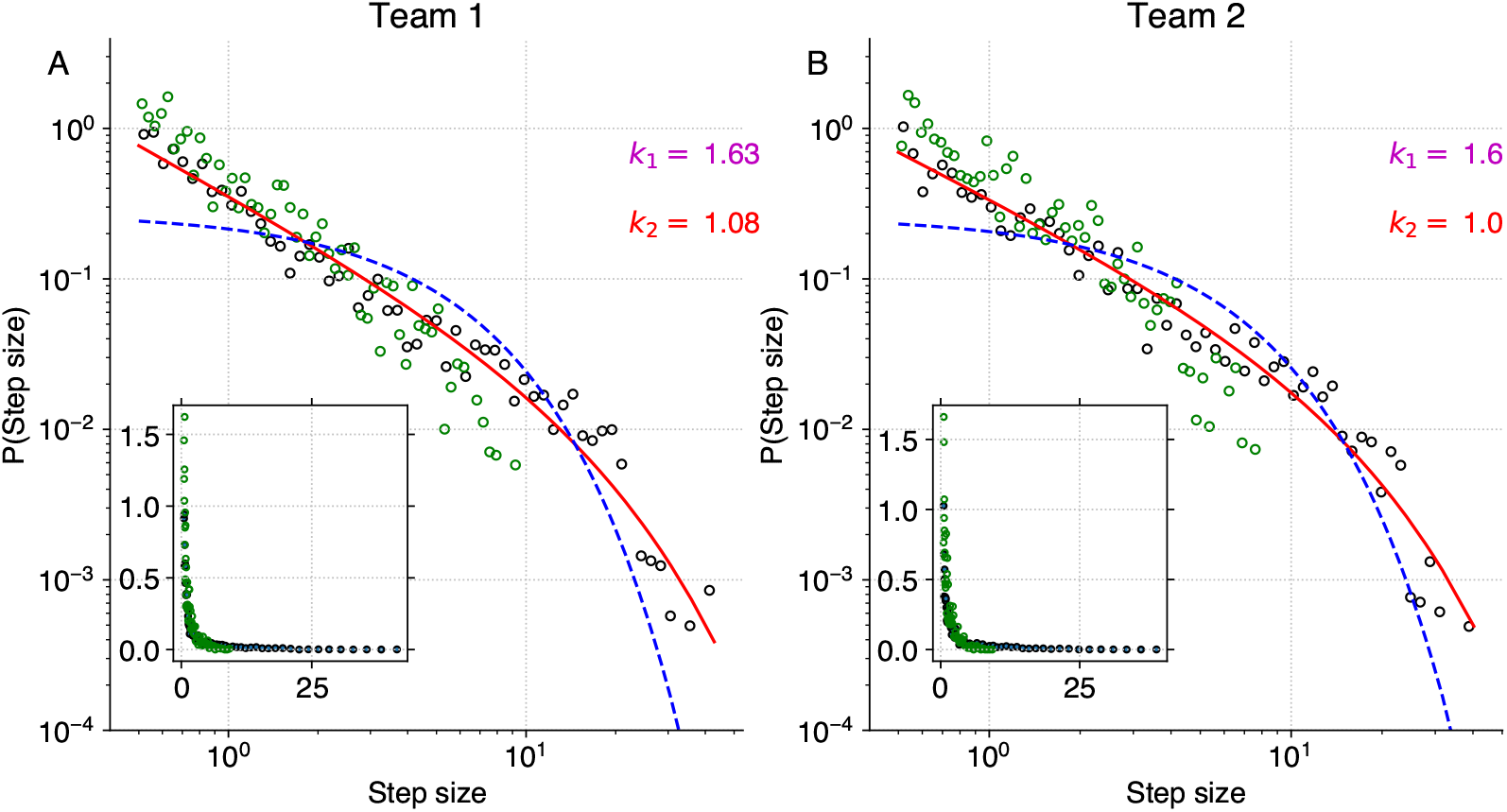
Step size distribution for centers’ of mass. Panes A and B depict the normalized distribution of step sizes for trajectories of the centers of mass of Teams 1 and 2, respectively. Black circles correspond to the real data, hollow circles represent synthetic data. The solid red line indicates the truncated power-law fit to the data. Power-law fit is omitted, however, its exponents *k*_1_ are given for both teams. Dashed blue line corresponds to the exponential fit to the real data. Insets show the the same data in the linear coordinates. Note the pronounced absence of the longer steps in the synthetic data.

### Analysis of multiple games

Additionally, we analyze all games contained in the 2022 dataset to elucidate if the Lévy walk exponent is in some way related to the player’s performance or his role in the team. Summarily, 643 individual trajectories of field players and 46 goalkeeper trajectories were analyzed, counting substitute players separately. To acquire insight into the game’s dynamics, for each player we compute two parameters: the mean distance of the player from the ball ⟨***D***_*ball*_⟩ and his mean distance from the center of mass of the team ⟨***D***_*team*_⟩. The former is meant to characterize players’ efficiency while the latter serves to quantify their level of involvement in team dynamics. Due to the distinct rules governing the goalkeepers’ movements, we choose to omit their data from the linear regression model, they are presented on the plots for the visual reference only.

Panes A and B in Figure 4 demonstrate that a statistically significant relationship exists between the power-law exponent of the Lévy walk distribution and the player’s distance to the ball, as well as with his distance to the center of his team (*p <* 0.05) in both cases. Pane C shows the relationship between the ⟨ ***D***_*ball*_ ⟩ and ⟨ ***D***_*team*_ ⟩. As the ball drives the dynamics of the football game this connection is to be expected. We have found that although the truncated power-law provides a better fit to the data, its exponent *k*_2_ has a lesser connection to the player’s dynamics, therefore *k*_1_, which corresponds to power-law fit without truncation, was used. Lower panes D, E, and F show the distribution of the ⟨ ***D***_*ball*_ ⟩, ⟨ ***D***_*team*_ ⟩, and *k*_1_ for all players. All said quantities are distributed normally, as the Kolmogorov-Smirnoff test for normality confirms.

**Figure 4.**
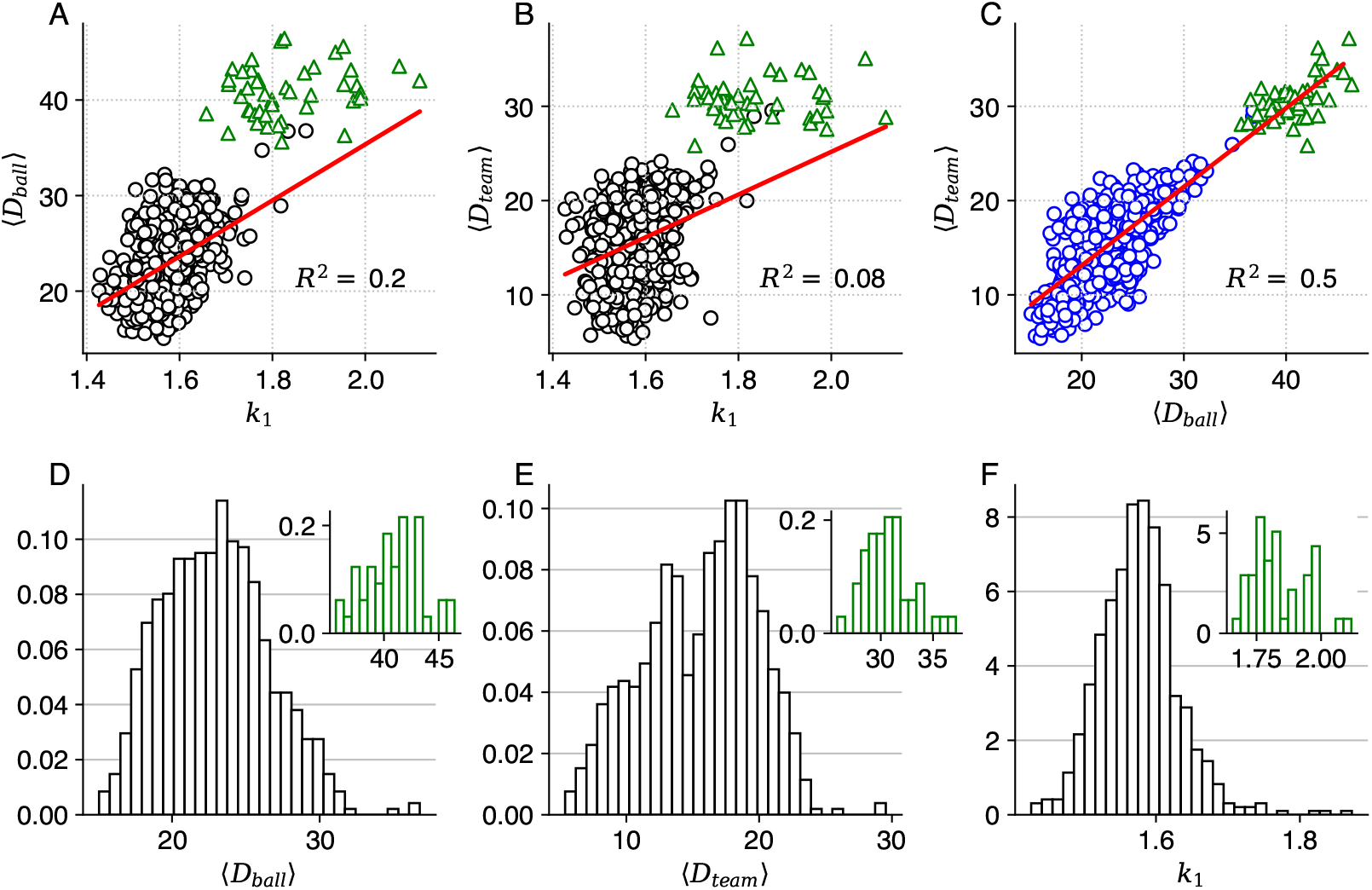
Lévy walk exponents and players’ performance. Pane A depicts the relationship between the *k*_1_ exponent of the step size distribution and mean distance to the ball ⟨***D***_*ball*_⟩, while pane B shows how players Lévy walk exponent is related to the average distance to the center of his team ⟨***D***_*team*_⟩ during the game. Pane C presents the relationship between the ⟨***D***_*ball*_⟩ and ⟨***D***_*team*_⟩. The regression slope is given with a solid red line. All distances are expressed in meters. Hollow green triangles correspond to the data of goalkeepers. Panes D, E, and F show the distributions (normalized histograms) of ⟨***D***_*ball*_⟩, ⟨***D***_*team*_⟩, and *k*_1_ for all players analyzed. Distributions of these quantities for goalkeepers are presented in the inset plots.

As gaining possession of the ball is a quintessential part of the football game we investigated if players behave differently when they have the ball, as compared to their movement otherwise during the game. As the properties of the step-size distribution do not depend on the specific ordering of elements it is possible to filter out the steps in which the player had an interaction with the ball and analyze them separately. We used trajectories of all field players to be found in the 2022 dataset and partitioned steps into two categories: steps in which the player had interaction with the ball and all remaining steps. These two aggregate distributions are presented in the Figure 5.

**Figure 5.**
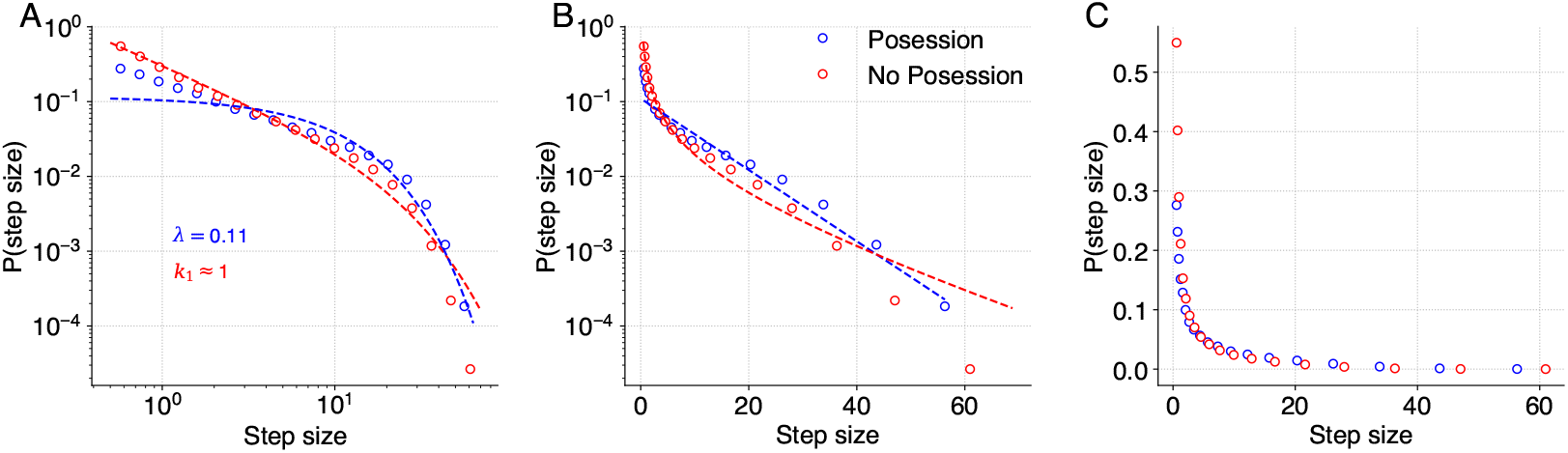
Steps size distribution departs from the power-law when players are in proximity of the ball. Aggregate step-size distribution for all field players from the dataset. Pane A presents the data in log-log coordinates while panes B and C show it on semi-logarithmic and linear axes. Blue circles correspond to the step-size distribution when the player is in possession of the ball, while red circles depict the steps when the player does not have the ball. Dashed blue and red lines correspond to max-likelihood estimates of the exponential and truncated power-law distributions respectively.

Max-likelihood methods have been used to fit several candidate distributions to the data. It should be noted that although the number of steps is vastly different between these two distributions in both cases overall the large number of data points allowed us to obtain confident max-likelihood estimates (*p >>* 0.05) of the best fit. When players are in possession of the ball the optimal candidate is the exponential distribution, however when players don’t have the ball truncated power-law is the best fit. It should be noted that this distinction is subtle and can only be inferred through log-likelihood tests of the aggregate distribution.

As discussed later Lévy walk exponent can be seen as an adaptive property for the organisms, therefore its slope could be influenced by both the player’s skill and their role in the team. For example, the goalkeepers’, whose movements are constrained by the rules the latter is the case.

At the same time, previous research indicates (***Cabrera and Milton, 2004***), that in human subjects the exponent changes as the subject becomes more proficient in the task. At this stage, we make no claims whether the observed relationship between the players’ metric and *k*_1_ is determined solely by their role in the team or if it is affected by their skill level as well.

## Discussion

Lévy walk-like behavior has been repetitively identified in human and animal locomotion. Although some of these findings have been challenged, due to improper data analysis most of them withstood the most rigorous statistical tests. To the best of our knowledge, Lévy walk has not yet been found in any of the sport’s practitioners, despite the fact the quantitative analysis of trajectories of professional players is becoming a common practice (***Sarmento et al., 2018***). Currently, the discourse surrounding the Lévy walk research is shifting from a mere debate regarding its plausibility in human and animal locomotion to a more nuanced discussion of its origin and purpose (***Reynolds, 2018***). Several compelling hypotheses exist to explain the underlying causes of the Lévy walk dynamics. In this work, we explore the conjecture that football games can be seen as a variation of the foraging behavior, as it has been suggested for human hunter-gatherers (***Reynolds et al., 2018***).

The game of football could be viewed as the type of foraging behavior in which agents seek to locate and engage the “resource” - the ball. This conjecture is supported by the observed relationship between the ⟨ ***D***_*ball*_ ⟩ and the power-law exponent *k*_1_. At the same time, football is a team endeavor in which individuals seek to coordinate their activity with their teammates and opponents. Some elements of the apparent coordination can be glimpsed from Figure 1, where we see periods of contraction and relaxation as well as evident interdependence between the trajectories of centroids of opposing teams. Intermittent dynamics of contraction and relaxation have been discovered in grazing sheep herd (***Ginelli et al., 2015***) where they serve to balance conflicting imperatives: optimal area coverage provided by the dispersed state and safety in numbers enabled by dense clustering. These oscillations could serve a similar role in teams’ dynamics during the game.

Deviation from the Lévy walk dynamics is observed when we separately consider the step-size distributions of players when they are in possession of the ball and otherwise (Figure 5). When players interact with the ball their aggregate step-size distribution is best described by the exponential, as is the case for Brownian motion, and in the second case, the truncated power-law distribution is recovered. It is tempting to interpret this result through the prism of the “foraging” approach to the game outlined above: agents resort to the Lévy behavior when they seek to locate and engage the resource, however when said resource is acquired impetus to forage disappears and movement patterns start to resemble Brownian motions. We would, however, be amiss not to mention that there might be a simpler mechanistic explanation to the observed phenomena if the need to interact with the ball constrains the player’s movements thereby making them resemble Brownian walkers. We should note, that although similarities can be found between the behavior of football players and that of foraging animals, one should be careful with that analogy as several added levels of behavioral complexity are present in a football game.

Inherent collective effects manifest themselves when we study the motion of the team’s centroids, which also exhibit Lévy walk dynamics. This finding is novel for Lévy walk research, as all previous discoveries concerned the motion of individuals. Transference of individual player’s movement patterns on the team level is facilitated by the high correlation between different team members. Said correlation is considered a hallmark of team behavior (***Marcelino et al., 2020***) in football and is informative of player performance and role. We interpret our finding as suggestive of the foraging activity performed not by a single individual but by a group, acting to maximize its collective performance. The “resource” foraged for, could be not a material object but a spatial configuration optimal to the needs of the moment. In our view, it would be instructive to investigate if similar phenomena could be observed in animals who practice collective hunting, such as wolves and hyenas.

One way to consider the broader significance of the uncovered phenomena is by invoking the work of Levin (***Levin, 2019, 2023***). One of his conjectures is that the impetus to understating the functioning of complex systems can be gained by expanding the definition of “Self”: leaving the idea of selfhood as something reserved to a single embodied and cognizant entity, such as a human or animal and entertaining the possibility of expanded or reduced selves, examples of which can be collectives or subsystems of an organism. “Selves” in such a paradigm should be treated as integrated entities capable of pursuing goals by modifying their behavior.

Once such a standpoint is taken concerning the Football team dynamics discovery of Lévy walk - a type of behavior that was previously only found in individuals on the collective level becomes something that could be anticipated. A football team is an entity that has a goal and can pursue it by modifying its behavior, both during the game and before it in the training process. It comes as no surprise then, that the movement strategy that the “Team self” adapts to optimize its performance shares statistical properties with the movement strategies adopted by individual “Selves” who experience similar environmental pressures.

## Conclusion

In our work, we showed that football players’ trajectories during the game manifest Lévy walk dynamics. Furthermore, Lévy walks exponents, which characterize the preponderance of longer steps in the distribution are related to the players’ role during the game, players with lower exponents are on average closer to the ball and explore a larger portion of the field. Curiously, a departure from the Lévy walk behavior is observed, when players are in possession of the ball.

Furthermore, power-law distribution of step sizes is present in the collective description of players’ activity (center of mass trajectory), an observation novel to the Lévy flight discourse, as it has been centered on the individual trajectory properties. At this stage we make no definitive claims regarding the generative mechanism of observed behavior, however, we theorize that the scale-free distribution of step sizes is influenced by both the collective nature of the game and its inherent “foraging” premise. Further work should include modeling to elucidate generative mechanisms as well as additional inquiry into the relationship between the Lévy walk exponent and other trajectory metrics and players’ performance. An alternative road, for other disciplines might be to study if the elements of foraging present in the football game might explain its widespread popularity.

## Methods and Materials

We obtained match recordings of Japan’s top football league, J-League, by licensing two datasets from the league’s official data company, DataStadium, Inc. Each dataset contains time-stamped trajectories of players from both teams for the sequence of games in which the season’s winning team participated. The data was acquired using the TRACAB’s optical video tracking systems Gen IV (***Linke et al., 2020***). In brief, this technology relies on gathering optical data from two multicamera units, located at both sides of the midfield line, which is then used to reconstruct players’ trajectories. Ball position is acquired in a separate process: when a player has the ball, his position is taken as a proxy for the ball’s coordinates, in between these events the position of the ball is obtained by linear interpolation.

The original data format is 25 Hz and the spatial resolution of the recording is in *cm*. Coordinates of all players from both teams were recorded, as well as the ball trajectory. To investigate if the temporal resolution of the data affects the results we have created several downsampled datasets with temporal resolution ranging from 1 Hz to 25 Hz and repeated our analysis. No statistically significant differences were observed between different downsamplings of the data.

All computations for the paper have been performed using custom scripts written in python programming language and are available from authors upon reasonable request. For fitting the power-law distribution to the data and performing log-likelihood *powerlaw* python package was used (***Alstott et al., 2014***). *x*_*min*_ value was set to 0.5 meters, to exclude potential spurious fluctuations. Steps with lengths below the threshold were excluded from the analysis. To investigate how the threshold value *x*_*min*_ affects the result we repeated our analysis using a range of threshold values: 0.5 *< x*_*min*_ *<* 5 *meters*. As it is expected the choice of threshold affects the value of the exponents *k*_1_ and *k*_2_ and once a sufficiently high threshold value is used the power-law distribution is reduced to the exponential. We have found that for most players’ when *x*_*min*_ ≤ 4 *meters* the power-law scaling is retained.

To filter out the steps in which the player had interacted with the ball, after partitioning the trajectory into steps using the 1D projection we computed the minimal distances between the player and the ball for each step separately. If said distance was ≤ 0.15 *meters* at least once, it was considered that a player had possession of the ball during that step. On average field players interact with the ball for 9% of their steps.

## Acknowledgments

This work was supported by JST Grant Number JPMJPF2205

## References

Alstott J, Bullmore E, Plenz D. powerlaw: a Python package for analysis of heavy-tailed distributions. PloS one. 2014; 9(1):e85777.

Berg HC. Random walks in biology. Princeton University Press; 1993.

Buldú JM, Busquets J, Echegoyen I, Seirul lo F. Defining a historic football team: Using Network Science to analyze Guardiola’s FC Barcelona. Scientific reports. 2019; 9(1):13602.

Cabrera JL, Milton JG. Human stick balancing: tuning Lévy flights to improve balance control. Chaos: An Interdisciplinary Journal of Nonlinear Science. 2004; 14(3):691–698.

Cotta C, Mora AM, Merelo JJ, Merelo-Molina C. A network analysis of the 2010 FIFA world cup champion team play. Journal of Systems Science and Complexity. 2013; 26(1):21–42.

Edwards AM, Phillips RA, Watkins NW, Freeman MP, Murphy EJ, Afanasyev V, Buldyrev SV, da Luz MG, Raposo EP, Stanley HE, et al. Revisiting Lévy flight search patterns of wandering albatrosses, bumblebees and deer. Nature. 2007; 449(7165):1044–1048.

Frencken W, Lemmink K, Delleman N, Visscher C. Oscillations of centroid position and surface area of soccer teams in small-sided games. European journal of sport science. 2011; 11(4):215–223.

Frencken W, Poel Hd, Visscher C, Lemmink K. Variability of inter-team distances associated with match events in elite-standard soccer. Journal of sports sciences. 2012; 30(12):1207–1213.

Ginelli F, Peruani F, Pillot MH, Chaté H, Theraulaz G, Bon R. Intermittent collective dynamics emerge from conflicting imperatives in sheep herds. Proceedings of the National Academy of Sciences. 2015; 112(41):12729– 12734.

Gonzalez MC, Hidalgo CA, Barabasi AL. Understanding individual human mobility patterns. nature. 2008; 453(7196):779–782.

Harris TH, Banigan EJ, Christian DA, Konradt C, Tait Wojno ED, Norose K, Wilson EH, John B, Weninger W, Luster AD, et al. Generalized Lévy walks and the role of chemokines in migration of effector CD8+ T cells. Nature. 2012; 486(7404):545–548.

Huizinga J. Homo ludens ils 86. Routledge; 2014.

Humphries NE, Weimerskirch H, Sims DW. A new approach for objective identification of turns and steps in organism movement data relevant to random walk modelling. Methods in Ecology and Evolution. 2013; 4(10):930–938.

de Jager M, Weissing FJ, Herman PM, Nolet BA, van de Koppel J. Lévy walks evolve through interaction between movement and environmental complexity. Science. 2011; 332(6037):1551–1553.

Levin M. The computational boundary of a “self”: developmental bioelectricity drives multicellularity and scale-free cognition. Frontiers in psychology. 2019; 10:493866.

Levin M. Darwin’s agential materials: evolutionary implications of multiscale competency in developmental biology. Cellular and Molecular Life Sciences. 2023; 80(6):142.

Linke D, Link D, Lames M. Football-specific validity of TRACAB’s optical video tracking systems. PloS one. 2020; 15(3):e0230179.

Marcelino R, Sampaio J, Amichay G, Gonçalves B, Couzin ID, Nagy M. Collective movement analysis reveals coordination tactics of team players in football matches. Chaos, Solitons & Fractals. 2020; 138:109831.

Michalet X. Mean square displacement analysis of single-particle trajectories with localization error: Brownian motion in an isotropic medium. Physical Review E. 2010; 82(4):041914.

Miramontes O, DeSouza O, Paiva LR, Marins A, Orozco S. Lévy flights and self-similar exploratory behaviour of termite workers: beyond model fitting. PloS one. 2014; 9(10):e111183.

Ramos-Fernández G, Mateos JL, Miramontes O, Cocho G, Larralde H, Ayala-Orozco B. Lévy walk patterns in the foraging movements of spider monkeys (Ateles geoffroyi). Behavioral ecology and Sociobiology. 2004; 55:223–230.

Reynolds AM, Ouellette NT. Swarm dynamics may give rise to Lévy flights. Scientific reports. 2016; 6(1):30515.

Reynolds A, Ceccon E, Baldauf C, Karina Medeiros T, Miramontes O. Lévy foraging patterns of rural humans. PlOS one. 2018; 13(6):e0199099.

Reynolds AM. Beating the odds in the aerial lottery: passive dispersers select conditions at takeoff that maximize their expected fitness on landing. The American Naturalist. 2013; 181(4):555–561.

Reynolds AM. Current status and future directions of Lévy walk research. Biology open. 2018; 7(1):bio030106.

Ríos-Uzeda B, Brigatti E, Vieira M. Lévy like patterns in the small-scale movements of marsupials in an unfamiliar and risky environment. Scientific reports. 2019; 9(1):2737.

Sarmento H, Clemente FM, Araújo D, Davids K, McRobert A, Figueiredo A. What performance analysts need to know about research trends in association football (2012–2016): A systematic review. Sports medicine. 2018; 48:799–836.

Sims DW, Humphries NE, Hu N, Medan V, Berni J. Optimal searching behaviour generated intrinsically by the central pattern generator for locomotion. Elife. 2019; 8:e50316.

Sims DW, Reynolds AM, Humphries NE, Southall EJ, Wearmouth VJ, Metcalfe B, Twitchett RJ. Hierarchical random walks in trace fossils and the origin of optimal search behavior. Proceedings of the National Academy of Sciences. 2014; 111(30):11073–11078.

Sims DW, Southall EJ, Humphries NE, Hays GC, Bradshaw CJ, Pitchford JW, James A, Ahmed MZ, Brierley AS, Hindell MA, et al. Scaling laws of marine predator search behaviour. Nature. 2008; 451(7182):1098–1102.

Stumpf MP, Porter MA. Critical truths about power laws. Science. 2012; 335(6069):665–666.

Viswanathan GM, Afanasyev V, Buldyrev SV, Murphy EJ, Prince PA, Stanley HE. Lévy flight search patterns of wandering albatrosses. Nature. 1996; 381(6581):413–415.

Viswanathan GM, Da Luz MG, Raposo EP, Stanley HE. The physics of foraging: an introduction to random searches and biological encounters. Cambridge University Press; 2011.

