## Supplementary materials for "Football as foraging? Movements by individual players and whole teams exhibit Lévy walk dynamics"

### Supplementary materials for the paper: "Football as foraging? Movements by individual players and whole teams exhibit Lévy walk dynamics"

This document contains supplementary information supporting the results of the main paper. Tables 1 and 2 provide additional information for all players, including the substitutes for both teams that participated in the game played on the 30th of July 2022. Detailed analysis of this match is contained in the **Analysis of a Single match** section of the main paper. For each player four estimates of the Lévy exponents $k$ are given: for X and Y dimensions, and power-law with truncation and without it. In all cases fitting was done using the maximum likelihood estimation, as described in the **Methods and Materials** section in the main body of the paper.

| **Player ID** | $k_1$ X | $k_2$ X | $k_1$ Y | $k_2$ Y |
|---|---|---|---|---|
| 1 | 1.66 | 1.08 | 1.88 | 1.43 |
| 2 | 1.46 | 1.00 | 1.61 | 1.08 |
| 4 | 1.44 | 1.00 | 1.61 | 1.09 |
| 5 | 1.46 | 1.00 | 1.60 | 1.19 |
| 7 | 1.42 | 1.00 | 1.65 | 1.19 |
| 8 | 1.45 | 1.00 | 1.60 | 1.00 |
| 10 | 1.50 | 1.00 | 1.62 | 1.15 |
| 11 | 1.46 | 1.00 | 1.61 | 1.10 |
| 14 | 1.45 | 1.00 | 1.58 | 1.03 |
| 23 | 1.45 | 1.00 | 1.61 | 1.06 |
| 24 | 1.43 | 1.00 | 1.57 | 1.04 |
| 27 | 1.45 | 1.00 | 1.57 | 1.02 |
| 32 | 1.46 | 1.00 | 1.62 | 1.07 |
| 45 | 1.47 | 1.00 | 1.65 | 1.22 |

**Table 1:** Additional information on Team 1.The following notation for Lévy exponents is used $k_1$ X - power-law for movements in X dimension, $k_1$ Y - powerlaw for movements in Y dimension, $k_2$ X -truncated power-law for movements in X dimension, $k_2$ Y - truncated powerlaw for movements in Y-dimension. Player IDs correspond to numbers on players' jerseys. All numbers are rounded to the second digit after the decimal.

| **Player ID** | $k_1$ X | $k_2$ X | $k_1$ Y | $k_2$ Y |
|---|---|---|---|---|
| 1 | 1.62 | 1.00 | 1.82 | 1.06 |
| 2 | 1.51 | 1.00 | 1.74 | 1.26 |
| 3 | 1.46 | 1.00 | 1.62 | 1.09 |
| 4 | 1.44 | 1.00 | 1.63 | 1.11 |
| 6 | 1.42 | 1.00 | 1.60 | 1.25 |
| 10 | 1.46 | 1.00 | 1.61 | 1.18 |
| 11 | 1.43 | 1.00 | 1.65 | 1.26 |
| 13 | 1.48 | 1.00 | 1.67 | 1.22 |
| 14 | 1.42 | 1.00 | 1.56 | 1.00 |
| 15 | 1.43 | 1.00 | 1.57 | 1.06 |
| 16 | 1.45 | 1.00 | 1.56 | 1.00 |
| 17 | 1.43 | 1.00 | 1.62 | 1.08 |
| 20 | 1.44 | 1.00 | 1.63 | 1.16 |
| 29 | 1.46 | 1.00 | 1.50 | 1.00 |

**Table 2:** Additional information on Team 2. Same notations is used as in Table 1

Table 3 gives a brief overview of the dataset which contains 2022 J-league matches in which the seasons' champion (Hokkaido Consadole Sapporo) participated. These games have been analyzed in the **Analysis of multiple games** section in the main body of the paper.

| **Home Team Name** | **Away Team Name** | **Date** | **Score** |
|---|---|---|---|
| Shimizu S-Pulse | Hokkaido Consadole Sapporo | 19/02/2022 | 1:1 |
| Hokkaido Consadole Sapporo | Sanfrecce Hiroshima | 26/02/2022 | 1:1 |
| Avispa Fukuoka | Hokkaido Consadole Sapporo | 06/03/2022 | 0:0 |
| Hokkaido Consadole Sapporo | Yokohama F Marinos | 12/03/2022 | 1:1 |
| Cerezo Osaka | Hokkaido Consadole Sapporo | 19/03/2022 | 2:2 |
| Hokkaido Consadole Sapporo | Urawa Reds | 02/04/2022 | 1:1 |
| Sagan Tosu | Hokkaido Consadole Sapporo | 06/04/2022 | 5:0 |
| Nagoya Grampus | Hokkaido Consadole Sapporo | 10/04/2022 | 0:2 |
| Hokkaido Consadole Sapporo | F.C.Tokyo | 16/04/2022 | 0:0 |
| Hokkaido Consadole Sapporo | Shonan Bellmare | 29/04/2022 | 1:0 |
| Gamba Osaka | Hokkaido Consadole Sapporo | 04/05/2022 | 0:0 |
| Hokkaido Consadole Sapporo | Kyoto Sanga F.C. | 07/05/2022 | 1:0 |
| Kashima Antlers | Hokkaido Consadole Sapporo | 14/05/2022 | 4:1 |
| Jubilo Iwata | Hokkaido Consadole Sapporo | 22/05/2022 | 1:2 |
| Hokkaido Consadole Sapporo | Kashiwa Reysol | 25/05/2022 | 1:6 |
| Vissel Kobe | Hokkaido Consadole Sapporo | 29/05/2022 | 4:1 |
| Kawasaki Frontale | Hokkaido Consadole Sapporo | 18/06/2022 | 5:2 |
| Hokkaido Consadole Sapporo | Gamba Osaka | 26/06/2022 | 1:0 |
| Kyoto Sanga F.C. | Hokkaido Consadole Sapporo | 02/07/2022 | 2:1 |
| F.C.Tokyo | Hokkaido Consadole Sapporo | 06/07/2022 | 3:0 |
| Hokkaido Consadole Sapporo | Kashima Antlers | 10/07/2022 | 0:0 |
| Kashiwa Reysol | Hokkaido Consadole Sapporo | 16/07/2022 | 1:0 |
| *Hokkaido Consadole Sapporo* | *Nagoya Grampus* | *30/07/2022* | *2:2* |
| Shonan Bellmare | Hokkaido Consadole Sapporo | 07/08/2022 | 1:5 |
| Hokkaido Consadole Sapporo | Vissel Kobe | 13/08/2022 | 0:2 |
| Hokkaido Consadole Sapporo | Sagan Tosu | 20/08/2022 | 1:2 |
| Hokkaido Consadole Sapporo | Cerezo Osaka | 02/09/2022 | 2:1 |
| Hokkaido Consadole Sapporo | Jubilo Iwata | 11/09/2022 | 4:0 |
| Yokohama F Marinos | Hokkaido Consadole Sapporo | 18/09/2022 | 0:0 |
| Hokkaido Consadole Sapporo | Kawasaki Frontale | 01/10/2022 | 4:3 |
| Hokkaido Consadole Sapporo | Avispa Fukuoka | 08/10/2022 | 1:2 |
| Urawa Reds | Hokkaido Consadole Sapporo | 12/10/2022 | 1:1 |
| Sanfrecce Hiroshima | Hokkaido Consadole Sapporo | 29/10/2022 | 1:2 |
| Hokkaido Consadole Sapporo | Shimizu S-Pulse | 05/11/2022 | 4:3 |

**Table 3:** Summary information regarding the games contained in 2022 dataset. Team names, dates and scores are provided. A single game which was analysed in depth in the paper is emphasized with italic.

#### Step size distribution dependence on players' position

To supplement the paper's main results, we investigate how the Lévy exponent of the step-size distribution is affected by players' position. We implement a modified version of methodology used to elucidate the influence of ball possession on players' dynamics. After compiling the aggregate step-length distribution for all field players in the dataset, we compute the maximum distance from the home edge of the field for each step. This distance distribution is shown in the pane B, Figure 1. Notably, the histogram implies that most movements occur in the mid-field zone, however prominent sub-peaks indicate enhanced activity in the vicinity of teams' own and opponents' field edges. Said observation stems from the dynamics of the football game in a manner obvious enough to necessitate no further elaboration.

Subsequently, we split the entire playing field into 8 rectangular parts in X-dimension, each approximately 13.1 meters in width. We use this partition to filter out the steps, which occur within the bounds of these zones. Note, that the field is partitioned in X-dimension, while step-lengths are computed in Y-projection, otherwise the results would have been trivial. We follow the same procedure for quantifying the goodness of fit for these 8 distributions and determining their parameters as described in **Methods and Materials**. In all cases, the truncated power-law emerges as the best fit for the data. However, Lévy exponents were found to be statistically different, when only steps that occurred in the immediate proximity of the home edge of the field were considered. Figure 1 pane B presents the aforementioned difference exhibiting two distributions, for the close vicinity of the home edge and opponent edge respectively. It can be inferred that smaller steps are more prominent when the players' are closer to their own side of the field.

#### Player's convex hull

Commonly used metric in the analysis of the spatially embedded data is the area of convex hull a minimal polygon that encompasses all points in the trajectory. Its area is informative of the portion of the field explored by each player during the game. Convex hulls are computed for each player, two separate values are computed for two halves of the game, and their average is used. The whole dataset of the year 2022 is used, as in the **Analysis of multiple games** section of the paper. Figure 2 demonstrates significant differences in the sizes of the players' convex hull areas. Goalkeepers retain their distinct status, as their hull areas are considerably smaller than those of the field players. To maintain consistency with the analysis in the main body of the paper goalkeepers were excluded from the linear model. A statistically significant relationship ($p<0.05$) exists between the area of the hull and the Lévy walk exponent, however, the percentage of explained variance is small. Furthermore, no statistically significant relationship exists between the hull area and the mean distance to the ball $\langle D \rangle$, when only field players are considered. Overall, the results imply that the metric is not sufficiently subtle to quantify the performance and role of the football players and is only weekly related to the Lévy exponent.

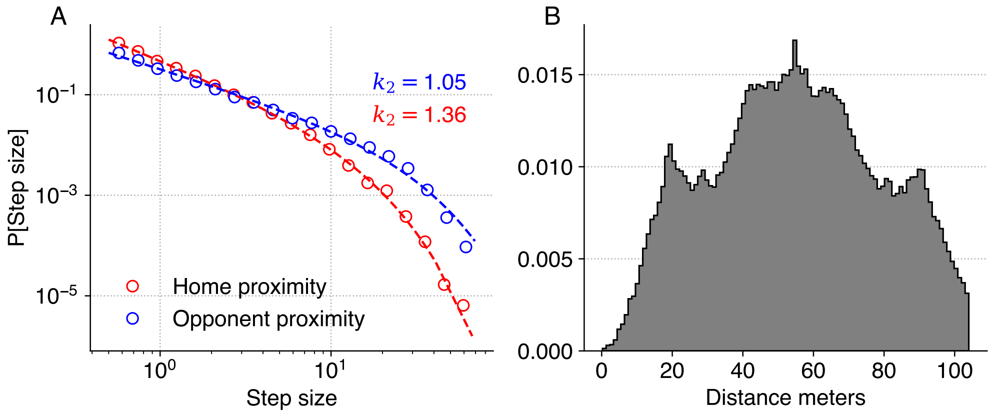


**Figure 1:** Players' location influences their movement patterns. Pane A: Log-binned distributions and truncated power-law fit for steps that occurred 13 meters away from the home edge (red circles) and 13 meters away from the opponent edge of the field (blue circles). Pane B: Normalized histogram of maximal distance from the home edge of the field for all players.

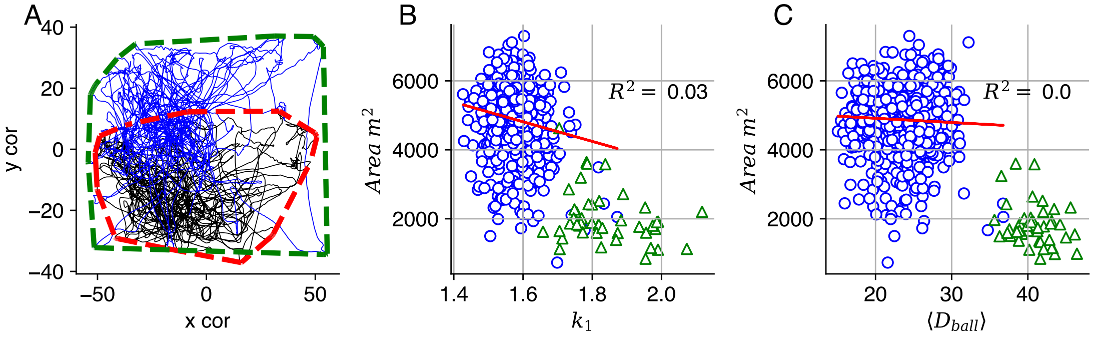


**Figure 2:** Lévy walks exponents and the area coverage. Pane A exhibits two illustrative trajectories for players from one team for the first half of the game. Red and green dashed lines show concave hulls encompassing the players' trajectory. The area of the green hull is $\approx 7304\ m^2$ and the red $\approx 3889\ m^2$. Lévy exponents $k_1$ computed for the whole duration of the game for these players are 1.6 and 1.61, respectively. Pane B presents the relationship between the players' hull and $k_1$ and pane C shows the relationship between the players' hull and his mean distance to the ball $\langle D_{ball} \rangle$. Solid red lines show linear regression fits, blue circles correspond to field players, and green triangles to goalkeepers.
